# Proteomic profiling identifies biomarkers of COVID-19 severity

**DOI:** 10.1101/2022.11.29.518411

**Authors:** Noa C. Harriott, Amy L. Ryan

## Abstract

SARS-CoV-2 infection remains a major public health concern, particularly for the aged and those individuals with co-morbidities at risk for developing severe COVID-19. Understanding the pathogenesis and biomarkers associated with responses to SARS-CoV-2 infection remain critical components in developing effective therapeutic approaches, especially in cases of severe and long-COVID-19. In this study blood plasma protein expression was compared in subjects with mild, moderate, and severe COVID-19 disease. Evaluation of an inflammatory protein panel confirms upregulation of proteins including TNFβ, IL-6, IL-8, IL-12, already associated with severe cytokine storm and progression to severe COVID-19. Importantly, we identify several proteins not yet associated with COVID-19 disease, including mesothelin (MSLN), that are expressed at significantly higher levels in severe COVID-19 subjects. In addition, we find a subset of markers associated with T-cell and dendritic cell responses to viral infection that are significantly higher in mild cases and decrease in expression as severity of COVID-19 increases, suggesting that an immediate and effective activation of T-cells is critical in modulating disease progression. Together, our findings identify new targets for further investigation as therapeutic approaches for the treatment of SARS-CoV-2 infection and prevention of complications of severe COVID-19.

## Introduction

Beginning in 2019, the global coronavirus disease (COVID-19) pandemic has presented a large-scale public health challenge, with the death toll in the United States exceeding 1 million (https://covid.cdc.gov/covid-data-tracker/#datatracker-home), and the global death count over 6 million as of July 2022 (https://covid19.who.int/). COVID-19, which results from infection by the severe acute respiratory syndrome coronavirus 2 (SARS-CoV-2) virus (Zhu et al., 2020), is a disease primarily characterized by dry cough, fever, and fatigue. However, symptoms can also include sore throat, shortness of breath, loss of smell and/or taste, headache, chills, nausea, vomiting, and diarrhea (Guan et al., 2020; Song et al., 2021; Wang et al., 2020a). Symptoms can also persist long after resolution of the initial infection, in some cases more than 14 months (de Miranda et al., 2022; Desai et al., 2022; Rass et al., 2022). In addition to this wide variety of symptoms, COVID-19 is associated with significant variation in disease severity. While the majority of cases are mild or asymptomatic (>85%), ~14% of cases require hospitalization, and <2% of all cases are lethal (Wu and McGoogan, 2020).

Clinical observations rapidly identified age as a primary risk factor for hospitalization and mortality (Buttenschon et al., 2022; Schafer et al., 2022). Age-related risk for severe COVID-19 has been a core focus of scientific investigation, and a variety of plausible explanations have been presented in the literature. Expression of angiotensin converting enzyme 2 (ACE2), the primary cell surface receptor for SARS-CoV-2 (Hoffmann et al., 2020; Letko et al., 2020), in addition to other SARS-CoV-2 entry factors, has been proposed as an explanation for age-related differences, with higher expression levels of these entry factors being detected in the nasal epithelium of older patients (Abrehart et al., 2022; Singh et al., 2020). In addition to the airway epithelium, ACE2 is also expressed on endothelial cells (Ferrario et al., 2005), where infection of the endothelial lining has been proposed to contribute to endotheliitis observed in severe COVID-19 (Varga et al., 2020). In general, children have comparatively healthier endothelium, potentially contributing to age-related differences in COVID-19 severity (Zimmermann and Curtis, 2022). Other studies, however, have found no correlation between proportion of ACE2+ cells and disease severity (Ma et al., 2021).

Building on these hypotheses, a more robust innate immune response in response to infection with SARS-CoV-2 is observed in children compared to adults (Loske et al., 2022; Pierce et al., 2021), and a dysregulation of innate immune function, similar to the “inflammaging” reported in aged populations, is observed in severe COVID-19 (Schultze and Aschenbrenner, 2021). Increased efficiency of the adaptive immune system is associated with children, and while adults may mount a more activated response, children are known to maintain higher levels of regulatory cells and memory T-cells (Cohen et al., 2021; Loske *et al*., 2022; Petrara et al., 2021). Increased activation of the adaptive immune system in this context appears to be detrimental and results in increased T-cell exhaustion. In this case, chronic T-cell activation results in the impaired function commonly observed in older patients, and correlates to COVID-19 disease severity (Blank et al., 2019; Palmer et al., 2021; Westmeier et al., 2020; Yang et al., 2020). This aberrant regulation of the immune response can also be maintained over time (Files et al., 2021), a finding with implications for the investigation of “long-haul COVID”. This study was designed to investigate the disparity in clinical severity of COVID-19, also considering the impact of aging and ethnicity, through proteomic profiling of patient-derived plasma samples collected in the Keck School of Medicine (KSoM) Biospecimen Repository for COVID-19 at the University of Southern California (USC), representing the spectrum of COVID-19 disease severity.

## Methods

### Study approval

The study was approved by the institutional review board (IRB) of the University of Southern California (USC: Protocol#: HS-20-00519).

### Patient recruitment

Patient plasma samples were collected between 1 May 2020 and 9 June 2021 from patients seen at the Keck Hospital, Verdugo Hills, and Los Angeles (LA) County Hospital and stored in the University of Southern California (USC) COVID-19 Biospecimen Repository. None of the subjects were vaccinated. Samples were not analyzed for SARS-CoV-2 variant. For this study, patients were assigned anonymized, coded IDs and were grouped according to the following cohort definitions: severe, indicating subjects who were admitted to the ICU for COVID-19 treatment; moderate, indicating subjects who were hospitalized for COVID-19 treatment but who were not admitted to the ICU; mild, indicating subjects who tested positive for SARS-CoV-2 but did not require hospitalization; and control, indicating subjects who tested negative for SARS-CoV-2 upon admission to the ICU for treatment of other severe diseases. Population demographics for these cohorts are summarized in **Table 1**. Participants were predominantly Hispanic/Latinx (69%), reflecting the demographics of donors available from the source biorepository (57.4% Hispanic/Latinx, https://sc-ctsi.org/about/covid-19-biorepository). The mean age of participants in this study was 56.1±1.58 years (**Supplemental Table 2**).

**Table 1:** Summary of study subject demographics. Data represents summarized data from a total of 70 subjects used in the current study. All data is expressed as a percentage of the total of subjects.

| Demographic |  | COVID-19 Positive |  |  | COVID-19 Negative | All Cohorts |
| --- | --- | --- | --- | --- | --- | --- |
|  |  | Severe<br>(N=22) | Moderate<br>(N=22) | Mild<br>(N=10) | (N=16) | (N=70) |
| Age group | <18 years | 0 (0%) | 0 (0%) | 0 (0%) | 0 (0%) | 0 (0%) |
|  | 19-35 years | 0 (0%) | 2 (3%) | 2 (3%) | 0 (0%) | 4 (6%) |
|  | 36-50 years | 3 (4%) | 5 (7%) | 4 (6%) | 5 (7%) | 17 (24%) |
|  | 51-65 years | 9 (13%) | 10 (14%) | 4 (6%) | 8 (11%) | 31 (44%) |
|  | >65 years | 10 (14%) | 5 (7%) | 0 (0%) | 3 (4%) | 18 (26%) |
| Sex | Female | 8 (11%) | 7 (10%) | 4 (6%) | 8 (11%) | 27 (39%) |
|  | Male | 14 (20%) | 15 (21%) | 6 (9%) | 8 (11%) | 43 (61%) |
| Race | White or Caucasian | 19 (27%) | 21 (30%) | 9 (13%) | 13 (19%) | 62 (89%) |
|  | Black or African American | 0 (0%) | 0 (0%) | 0 (0%) | 1 (1%) | 1 (1%) |
|  | Asian | 1 (1%) | 0 (0%) | 1 (1%) | 1 (1%) | 3 (4%) |
|  | American Indian or Alaska Native | 0 (0%) | 1 (1%) | 0 (0%) | 0 (0%) | 1 (1%) |
|  | Native Hawaiian or other Pacific Islander | 0 (0%) | 0 (0%) | 0 (0%) | 0 (0%) | 0 (0%) |
|  | Other race | 0 (0%) | 0 (0%) | 0 (0%) | 0 (0%) | 0 (0%) |
|  | Unknown race | 2 (3%) | 0 (0%) | 0 (0%) | 1 (1%) | 3 (4%) |
| Ethnicity | Hispanic or Latinx | 18 (26%) | 17 (24%) | 5 (7%) | 8 (11%) | 48 (69%) |
|  | Not Hispanic or Latinx | 4 (6%) | 5 (7%) | 5 (7%) | 7 (10%) | 21 (30%) |
|  | Unknown ethnicity | 0 (0%) | 0 (0%) | 0 (0%) | 1 (1%) | 1 (1%) |

### Immunophenotyping

Plasma samples were analyzed for protein expression by Olink proximity extension assays (PEA) for quantification of 184 secreted markers. Olink’s Target 96 Inflammation and Target 96 Oncology II panels were chosen for the spread of proteins related to immune response. Each panel consisted of 92 proteins each, of which 6 proteins were included on both panels, resulting in 184 proteins in total and 178 unique proteins. In total, 144 samples were analyzed. Samples were determined to fail quality control if internal incubation and detection controls deviated +/-0.3 Normalized Protein eXpression (NPX) value from the median value across all samples. Four samples from the severe cohort, failed both panels and was excluded. Eight samples, one from the control cohort, four from the moderate cohort, and three from the severe cohort, failed the Oncology II panel and were excluded from analysis of that panel but were included in the analysis of the Inflammation panel. Seven proteins had NPX values under the protein-specific limit of detection (LOD) in >50% of samples in all cohorts, and were excluded from statistical analysis, leaving 171 unique proteins. The proteins excluded from analysis were IL-2RB, IL-1α, IL-2, β-NGF, IL-13, IL-33, and IL-4.

### Statistics

NPX data was processed using the platform-specific Olink Insights Stats Analysis app (https://olinkproteomics.shinyapps.io/OlinkInsightsStatAnalysis/). This app was used to perform platform-standard pairwise comparisons between cohorts using unpaired Student’s *t*-tests. These were performed for each individual protein included in the analysis panels. The Insights Stats Analysis app was also used to adjust p-values resulting from this analysis for multiple testing using the Benjamini-Hochberg method. False discovery rate (FDR) threshold was set to 0.05. Network analysis was performed using the STRING database (STRING Consortium, version 11.5). Protein-protein connections were assigned a combined “score” by evaluating probabilities of interaction derived from literature and database mining, then mapped according to these scores; full description of STRING analysis is described in (von Mering et al., 2005).

## Results

### Subject demographics and assay quality control

Blood plasma samples were obtained from the USC COVID-19 Biorepository and were collected from subjects seen at the Keck Hospital (52.9%), Verdugo Hills (12.9%) and Los Angeles County Hospital (34.3%) between 5/1/2020 and 6/9/2022. **Table 1** provides the core demographics of the subject population that provided samples for this project. The population of biobank donors was predominantly Hispanic/Latinx (57.4%, https://sc-ctsi.org/about/covid-19-biorepository), with samples unevenly distributed across all categories of COVID-19 severity. In cases requiring hospitalization, >75% of samples were from Hispanic/Latinx subjects. Subjects were segregated into four independent cohorts based on the hospitalization status of the patient. Categories of severe, moderate, mild and control were based on the following cohorts: 1) severe were COVID-19 positive subjects in the intensive care unit (ICU) being treated for COVID-related illness, 2) moderate were COVID-19 positive subjects that were hospitalized but not requiring ICU treatment, 3) mild were COVID-19 positive subjects that did not require hospitalization and 4) control were COVID-19 negative subjects that were treated in the ICU for other severe illness. The mean age of participants in the study across all categories was 56.1 ±1.58 years. 26% of the subjects were over 65 years and 6% were under 35 years. The mean age within each subject cohort is included in **Supplemental Table S1**. Overall, 61% of the subjects were male, 89% were White/Caucasian, and 69% were Hispanic/Latinx. For hospitalized COVID-19 patients in the severe and moderate groups, samples were obtained at day of admission (Day 1), Day 3, Day 5, and Day 7, where available. For the control cohort and mild cohort, the only sample evaluated was day of test/admission (Day 1).

Proteomic analysis of the plasma samples was completed using Olink^®^ proximity extension assays (PEA) for quantification of 184 secreted immunoregulatory biomarkers. The Olink Target 96 Inflammation and Target 96 Oncology II panels were chosen for the spread of proteins related to immune response. Each panel consisted of 92 proteins each, of which 6 proteins were analyzed on both panels, resulting in 184 proteins in total and 178 unique proteins. Samples were determined to fail quality control if internal incubation and detection controls deviated +/-0.3 Normalized Protein eXpression (NPX) value from the median value across all samples. Four samples, from the severe cohort, failed both panels and was excluded. Eight samples, one from the control cohort, four from the moderate cohort, and three from the severe cohort, failed only the Oncology II panel and were excluded from analysis of that panel but were included in the analysis of the Inflammation panel. Seven proteins had NPX values under the protein-specific limit of detection (LOD) in >50% of samples in all cohorts, and were excluded from downstream analysis, leaving 171 unique proteins.

### Plasma protein expression signatures are associated with severity of COVID-19

We first performed a principal component analysis (PCA) to highlight variation between proteomic signatures of plasma samples collected at Day 1 (**Figure 1A**). Samples in the severe COVID-19 cohort (**Figure 1A, red**) and moderate cohort (**Figure 1A, yellow**) cluster together, but are distinct from mild (**Figure 1A, blue**) and control cohorts (**Figure 1A, cyan**). Unbiased clustering by protein NPX values is visualized in the heatmap featured in **Figure 1B**. Again, the samples clustered by severity, indicating that protein expression signatures exist representing COVID-19 severity in our patient cohorts. To specifically evaluate changes in protein expression and their association with COVID-19 disease severity, pairwise comparisons of the mean NPX values for each protein were used to determine differentially expressed proteins (DEPs) between subject cohorts. Of the 171 unique proteins analyzed, 109 DEPs were observed between our study cohorts. DEPs between each study cohort are summarized in **Table 2**, and a comprehensive list of the significant DEPs across all study cohorts, as determined by unpaired Student’s *t*-tests of NPX value by cohort, is available in the full data set.

**Figure 1:**
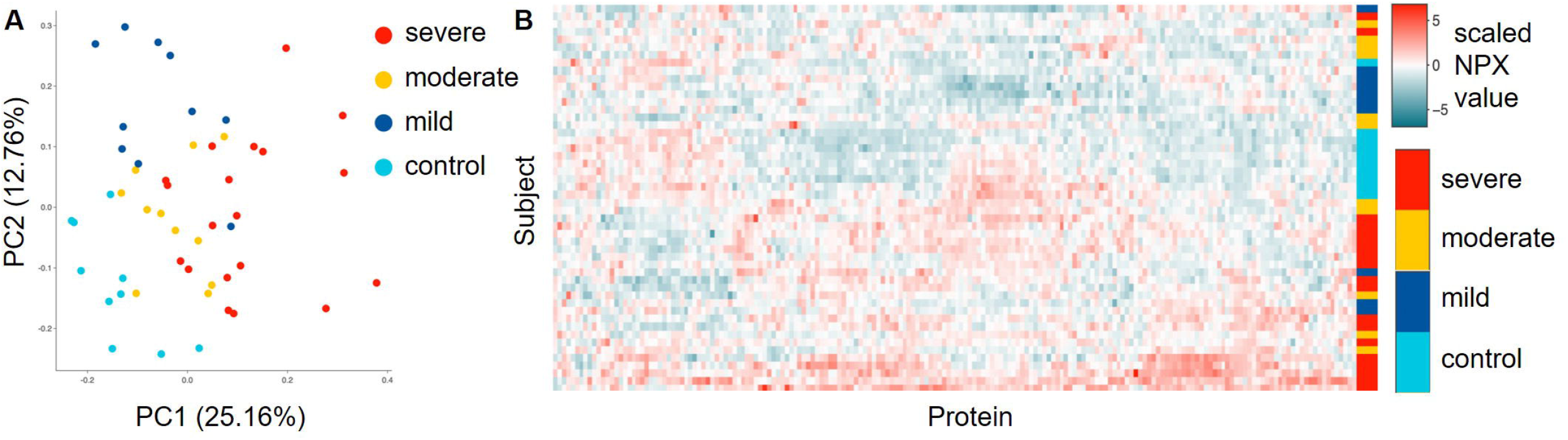
Plasma protein expression signatures are associated with severity of COVID-19. A) Principal component analysis (PCA) plot comparing samples across all cohorts. B) Heatmap of relative protein expression. Proteins were evaluated through an unbiased hierarchical clustering according to scaled NPX value for each sample.

**Table 2:** Summary of top DEPs comparing all subject cohorts. For each pairwise comparison, total DE indicates the total number of differentially expressed proteins (DEPs); up indicates the number of DEPs in that comparison that were increased, down indicates the number of DEPs in that comparison that were decreased. “Most sig” indicates the most significant DEP in each comparison; % change, p-value, and adjusted (adj.) p-value are listed for each DEP.

| Analysis | Total DE | Up | Down | Most sig. | % change | p-value | Adj. p-value |
| --- | --- | --- | --- | --- | --- | --- | --- |
| Severe vs mild | 109 | 85 | 24 | EN-RAGE | 479.03 | 4.96E-11 | 3.86E-09 |
| Severe vs moderate | 43 | 37 | 6 | TNFB | -40.16 | 1.57E-04 | 0.0213 |
| Severe vs control | 84 | 77 | 7 | MSLN | 258.83 | 3.69E-08 | 6.79E-06 |
| Mild vs moderate | 65 | 12 | 53 | IL6 | -85.88 | 7.46E-07 | 6.82E-05 |
| Mild vs control | 87 | 46 | 41 | TRANCE | 155.67 | 1.97E-07 | 3.63E-05 |
| Moderate vs control | 54 | 52 | 2 | GZMB | 178.31 | 7.17E-05 | 0.0049 |

### Network analysis identifies markers of inflammation and cell proliferation in severe COVID-19

To establish the identity of the proteomic signatures associated with the severity of a subject’s response to SARS-CoV-2 infection, we further compared DEPs between subject cohorts. To evaluate the connections within the biological processes associated with the DEPs in our targeted proteomics analysis, we performed STRING network analysis and Ingenuity Pathway Analysis (IPA) focusing specifically on the DEPs between severe COVID-19 and all other cohorts. As expected, two major “hubs” of protein interactions were observed centered around the core of the inflammation and oncology 2 proteomics panels. The inflammation hub is associated with the activation of both Th1 and pro-inflammatory cytokines and chemokines including tumor necrosis factor beta (TNFβ), interleukin-6 (IL-6), interleukin-8 (IL-8), interleukin-12 (IL-12), CXC motif chemokine ligand 9 (CXCL9), and CC motif chemokine ligand 3 (CCL3), as well as Th2 and anti-inflammatory cytokines and chemokines including interleukin-10 (IL-10), thymic stromal lymphopoietin (TSLP), and CC motif chemokine ligand 20 (CCL20) (**Figure 2A, red cloud**). Interleukin-17 (IL-17) signaling and Th1 and Th2 activation pathways were among the most significantly changed pathways specific to severe COVID-19 (**Figure 2B**). The “oncology 2” hub clustered around growth factors and growth factor receptors including fibroblast growth factor 5 (FGF-5), colony stimulating factor (CSF), ephrin type-A receptor 2 (EPHA2), protransforming growth factor alpha (TGF-α), and beta-nerve growth factor (β-NGF), suggesting significant activation of tissue regeneration, differentiation and survival signaling pathways (**Figure 2A, green cloud**). IPA also identified wound healing and airway pathologies associated with chronic lung disease as some of the most significantly changed pathways specific to severe COVID-19 (**Figure 2B**). In conjunction with tissue remodeling, we also identified connections between Fas-associated death domain protein (FADD), Fas ligand (FASLG), TNF-related apoptosis-inducing ligand (TRAIL), and caspase 8 (CASP-8), markers associated with apoptosis. IPA also validated the upregulation of a pathogenic response including granulocyte and agranulocyte adhesion, diapedesis, and pattern recognition receptor pathways (**Figure 2B**).

**Figure 2:**
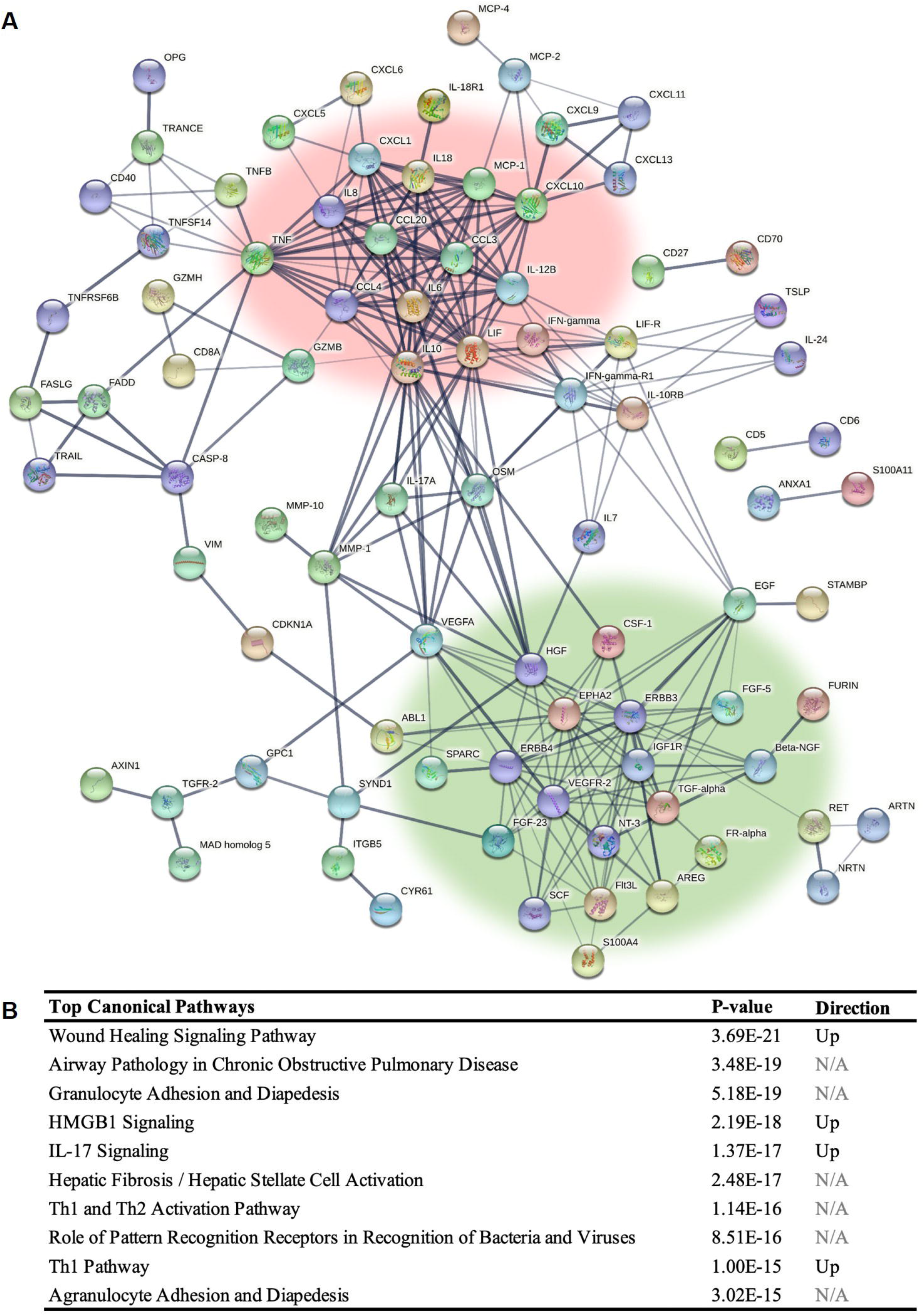
Network analysis identifies markers of inflammation and cell proliferation in severe COVID-19. A) STRING network analysis map of all significantly differentially expressed proteins (DEPs) in severe vs any other cohort. Lines indicate known associations between proteins. Thickness of line indicates confidence score (minimum = 0.4). Networks of associated markers are highlighted by background clouds; inflammatory markers in red, proliferative markers in green. B) Top 10 most enriched pathways as determined by Ingenuity Pathway Analysis (IPA) of all significant DEPs in severe vs any other cohort.

### Markers of inflammation and cell proliferation are expressed significantly higher in severe COVID-19

Volcano plots highlighting significant DEPs between the severe COVID-19 cohort and each of the other subject cohorts show 109 DEPs comparing the severe and mild COVID-19 cohorts (**Figure 3A**), 43 DEPs comparing the severe and moderate COVID-19 cohorts (**Figure 3B**), and 84 DEPs comparing the severe and control cohorts (**Figure 3C**). To determine a protein signature specific to severe COVID-19, DEPs between all paired analyses represented in the volcano plots were overlaid in a Venn diagram (**Figure 3D**). In total, 29 DEPs were significant in all comparisons (**Table 3**). Plots representing the samples included in each cohort highlight a severity-associated decline in the amount of plasma proteins detected, as shown for syndecan-1 (SYND1, **Figure 3E**), EN-RAGE (S100A12, **Figure 3F**), and mesothelin (MSLN, **Figure 3G**).

**Figure 3:**
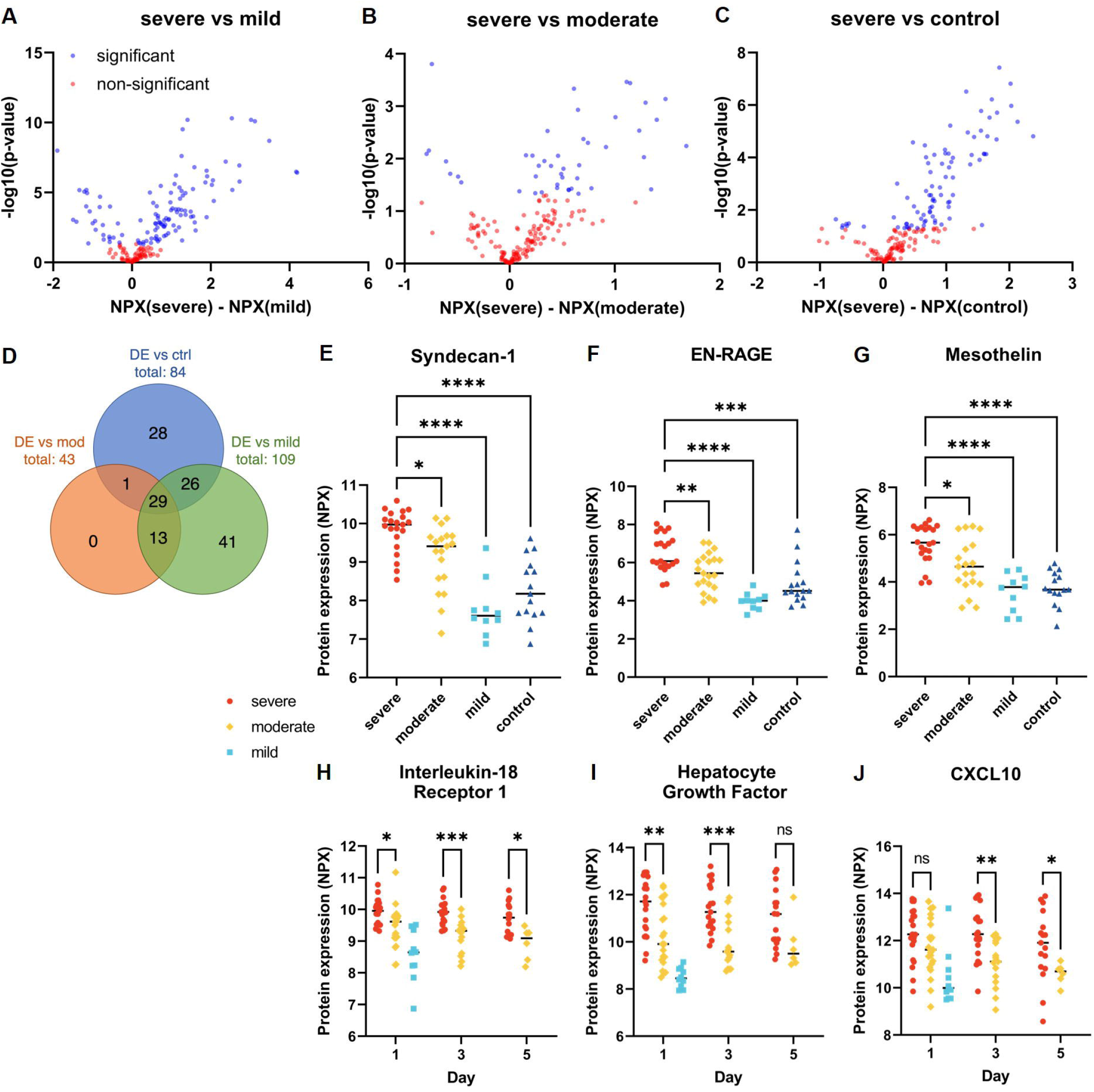
Markers of inflammation and cell proliferation are expressed significantly higher in severe COVID-19. A-C) Volcano plots showing DEPs between A) severe and mild, B) severe and moderate, and C) severe and control cohorts. Significance was determined by p<0.05. D) Venn diagram overlaying DEPs between severe and mild (green), severe and moderate (orange), and severe and control cohorts (blue). E-G) Cluster plots of normalized protein expression (NPX) values by cohort. E) Syndecan-1, F) EN-RAGE, and G) Mesothelin. H-J) Cluster plots of NPX values in severe and moderate cohorts over collection days 1, 3, and 5. Mild values at day 1 included for reference. H) Interleukin-18 Receptor 1, I) Hepatocyte Growth Factor, and J) CXCL10. For all cluster plots data is presented as: *p<0.05, **p<0.01, ***p<0.001, ****p<0.0001. Black horizontal line indicates the median.

**Table 3:**
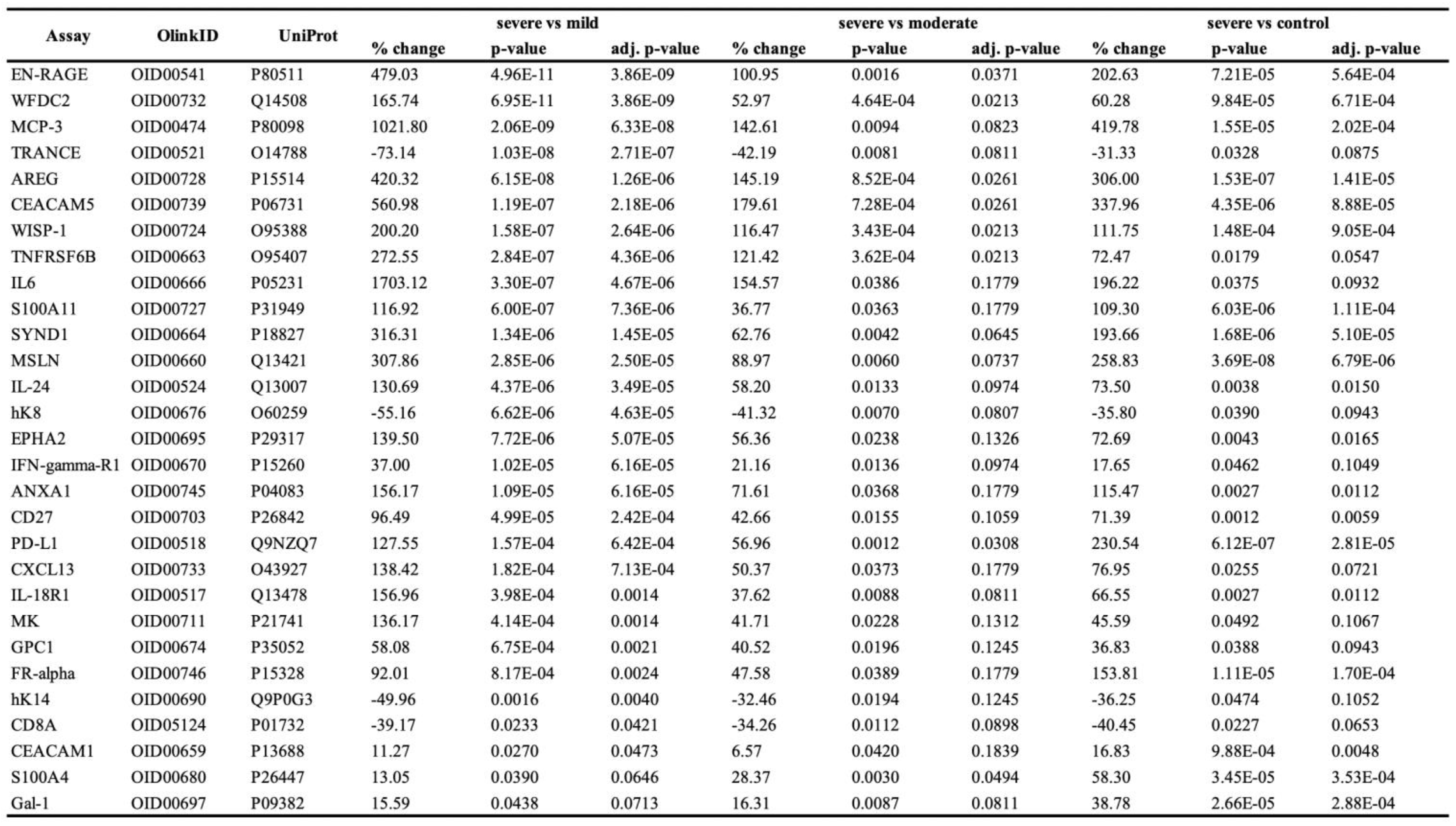
DEPs significant to severe COVID-19 versus all other cohorts. List of DEPs that were significant when comparing the severe cohort (N=22) to mild (N=10), moderate (N=22), and control (N=16) cohorts. % change, p-value, and adjusted (adj.) p-value are listed for each pairwise comparison of each DEP.

Interestingly, several proteins that were elevated in both severe and moderate COVID-19 on Day 1 differentially resolved over time in the moderate COVID-19 cohort, lowering to levels comparable to the mild cohort by Day 5 while remaining elevated in the severe COVID-19 cohort. These proteins included interleukin-18 receptor 1 (IL18-R1, **Figure 3H**), hepatocyte growth factor (HGF, **Figure 3I**), and CXC motif chemokine ligand 10 (CXCL10, **Figure 3J**). Such changes in the signaling pathways associated with these proteins likely underly the timely resolution of COVID-19 disease.

Ethnicity was established as a risk factor for severe COVID-19 early during the pandemic, following the observation of higher levels of hospitalization and more severe disease outcomes for Hispanic compared to non-Hispanic subjects (Chiumento et al., 2022; Renelus et al., 2021). Given the ethnic disparity in the Los Angeles community, the samples in our cohort were majority Hispanic/LatinX (69%, **Table 1**), providing an opportunity to evaluate the intersection of ethnicity and severity as characterized by proteomic profiling. 22 significant DEPs were identified between Hispanic and non-Hispanic subjects (**Supplemental Table S2**). All 22 of these DEPs were also significantly differentially expressed between severe cohort and at least one other cohort. 21 were significant between severe and mild cohorts, 18 between severe and control cohorts and 14 between severe and moderate cohorts. Disparity in disease outcome has also been associated with sex as a biological variable with males overrepresented in cases having severe complications from COVID-19 (Mikami et al., 2021). Our subject population comprised of 61% male and 39% female subjects. While the control cohort was evenly split, with 50% male and 50% female subjects, all COVID-19 cohorts were predominantly male (severe 64%, moderate 68% and mild 60%). Comparing NPX values between male and female subjects we observed no significant DEPs.

### A subset of “protective” proteins are significantly higher in mild disease also correlate to younger age groups

In search of biomarkers potentially associated with an efficient and “protected” response to SARS-CoV-2 infection DEPs unique to the mild COVID-19 cohort were identified. In total, 109 significant DEPs were identified comparing the mild to severe COVID-19 cohorts (**Figure 4A**), 65 comparing the mild to moderate COVID-19 cohorts (**Figure 4B**), and 87 comparing the mild to control cohorts (**Figure 4C**). Overlaying these DEPs in the Venn diagram in **Figure 4D** highlights 40 proteins specific to mild COVID-19 (**Table 4**). Most of these proteins were detected at significantly lower abundance in the mild cohort (80%, **Table 4**), likely representative of a tempered immune response and decreased persistence of pro-inflammatory cytokines and chemokines. DEPs that were significantly augmented in the mild subject cohort and downregulated with severity of COVID-19 included TNF superfamily member 11 (TRANCE aka RANK-L, **Figure 4E**), FASLG (**Figure 4F**), XPNPEP2 (**Figure 4G**), and CD207 (**Figure 4H**), suggesting expression of these proteins may be associated with a stronger response to fighting infection with SARS-CoV-2. In addition, osteoprotegerin (OPG), a decoy receptor for TRANCE, was more highly expressed in the severe cohort (**Supplemental Figure S1**). Interestingly, two of these proteins (TRANCE and FASLG) were also expressed at higher levels in the youngest patient cohort (19 to 35 years) decreasing in expression with age (**Figure 4I and 4J**). This finding is consistent with age being a predominant co-morbidity associated with COVID-19 (Mikami *et al*., 2021; Suleyman et al., 2022) and suggests that circulating levels of TRANCE and FASLG may be associated with a more effective response to SARS-CoV-2 infection.

**Figure 4:**
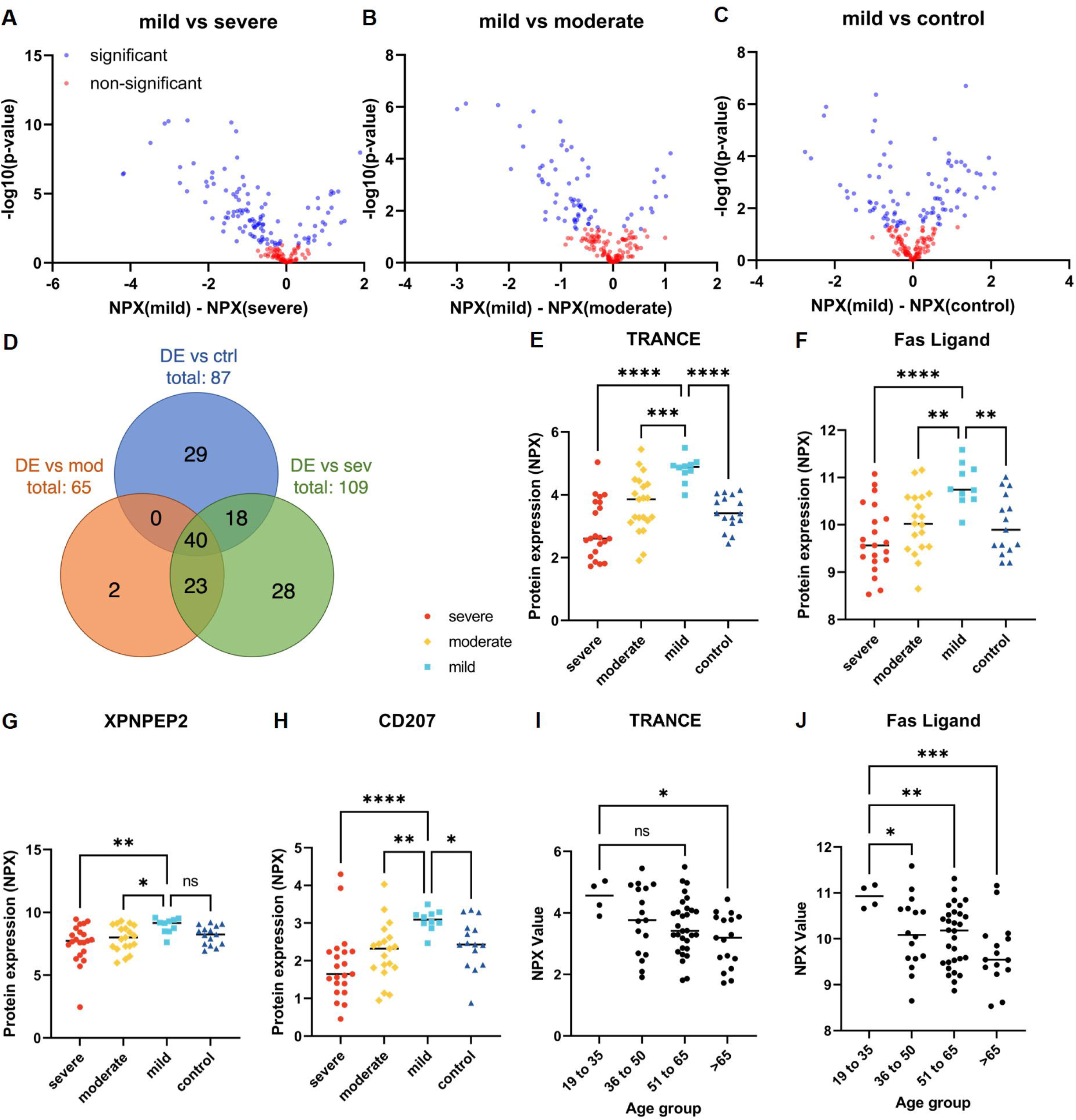
A subset of “protective” proteins are significantly higher in mild disease also correlate to younger age groups. A-C) Volcano plots showing DEPs between A) mild and severe, B) mild and moderate, and C) mild and control cohorts. Significance was determined by p<0.05. D) Venn diagram overlaying DEPs between mild and severe (green), mild and moderate (orange), and mild and control cohorts (blue). E-I) Cluster plots of normalized protein expression (NPX) values by cohort. E) TRANCE, F) Fas Ligand, G) XPNPEP2, and H) CD207. I-J) Cluster plots of NPX values by age group for I) TRANCE and J) Fas Ligand. For all cluster plots: *p<0.05, **p<0.01, ***p<0.001, ****p<0.0001. Black horizontal line indicates the median.

**Table 4:**
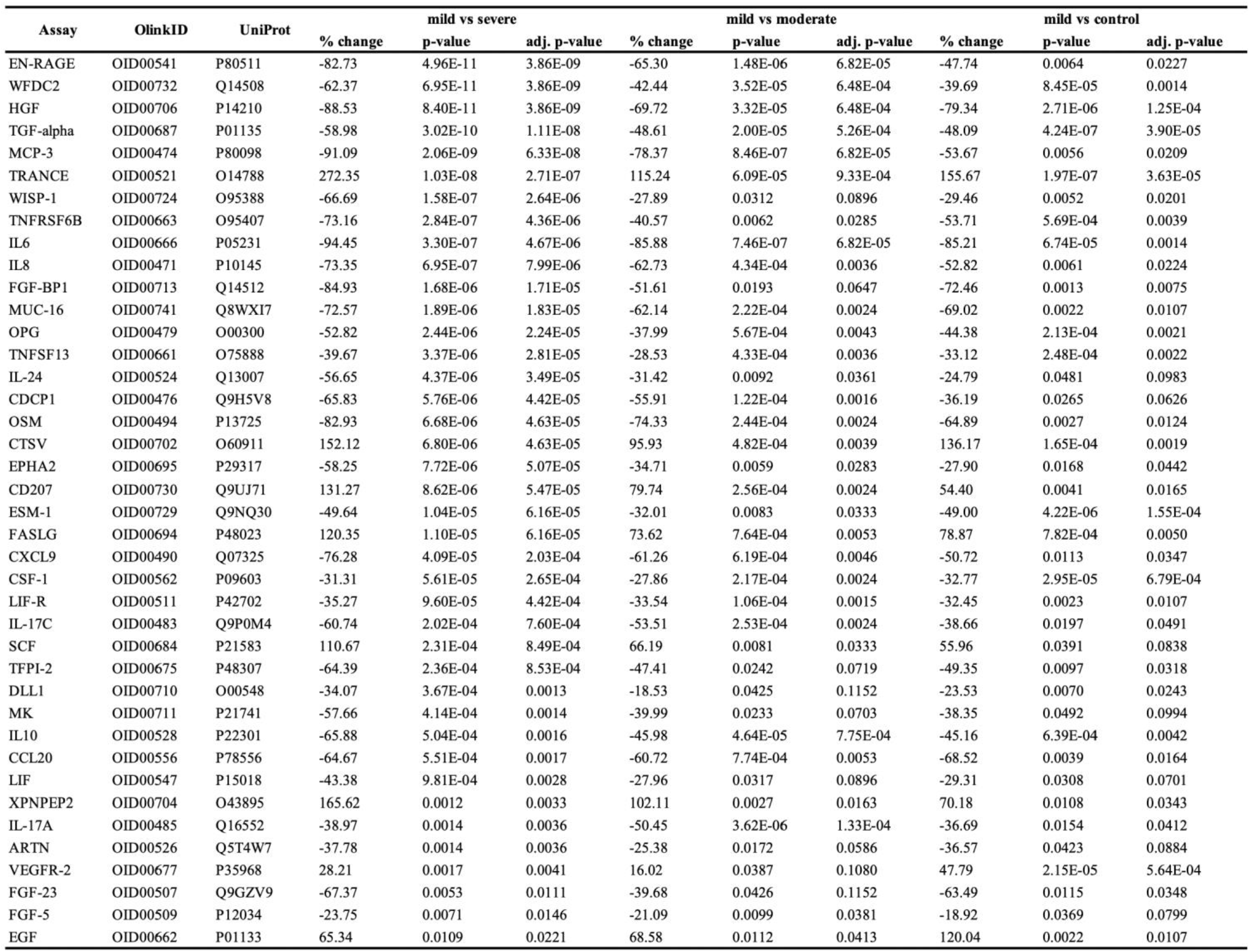
DEPs significant to mild COVID-19 versus every other cohort. List of DEPs that were significant when comparing the mild cohort (N=10) to severe (N=22), moderate (N=22), and control (N=16) cohorts. % change, p-value, and adjusted (adj.) p-value are listed for each pairwise comparison of each DEP.

## Discussion

One of the major challenges in managing the COVID-19 pandemic is the wide variation in disease severity, ranging from mild respiratory symptoms that resolve with minimal outpatient treatment to acute respiratory distress syndrome (ARDS) requiring ICU admission and mechanical ventilation. The phenomena underlying this disparity in disease progression are still poorly understood. In this study, we present data evaluating plasma protein expression that characterizes COVID-19 disease response by severity, evaluating proteins beyond the typical analysis of inflammation associated proteins. The signatures identified in our study include several proteins that have been previously associated with severe COVID-19 in both clinical datasets and *in vitro* models, as well as several new markers that have previously not been associated with the severity of response to SARS-CoV-2 infection. In severe COVID-19, the most significant and highly enriched proteins were associated with an inflammatory response to viral infection, including IL-6, IL-8, CCL3, CXCL9, and CXCL10. Several of these have ongoing clinical trials evaluating the effectiveness of regulating these proteins in preventing progression to severe COVID-19 and associated ARDS. Furthermore, our data correlates with previous findings showing a COVID-19-specific elevation of inflammatory signaling pathways (Song et al., 2020a). SYND1 and EN-RAGE, markers associated with COVID-19 disease severity and mortality (Maldonado et al., 2022; Russell et al., 2022; Zeng et al., 2021; Zhang et al., 2021) were among the most significantly elevated markers in our severe cohort.

SYND1 is a heparan sulfate proteoglycan, involved in leukocyte recruitment, vascular repair, and tumor angiogenesis (Teng et al., 2012). Increased expression of this marker is associated with acute endothelial glycocalyx degradation, as well as higher oxygen and mechanical life support requirements and mortality from COVID-19 (Dupont et al., 2021; Johansson et al., 2011; Karampoor et al., 2021; Ogawa et al., 2021). In addition to its significance as a potent marker of endothelial damage during the pathogenesis of COVID-19 (Stahl et al., 2020), both facilitation of viral entry via ACE2 co-localization (Hudak et al., 2021) and transmission of virus to epithelial cells via binding with dendritic cells (Bermejo-Jambrina et al., 2021) have been proposed as mechanisms by which SYND1 actively modulates SARS-CoV-2 infection, making it an important candidate as both a biomarker and a potential target for therapeutic intervention.

EN-RAGE is a pro-inflammatory calcium-binding protein associated with a variety of inflammatory conditions across organ systems, including rheumatoid and psoriatic arthritis, coronary heart disease, cystic fibrosis, autoimmune hepatitis, and Kawasaki disease (Foell et al., 2003a; Foell et al., 2003b; Foell et al., 2003c; Ligthart et al., 2014; Wu et al., 2020). Elevated levels of EN-RAGE have been shown to correlate to increased inflammation in response to COVID-19 (Thwaites et al., 2021), including in asymptomatic cases up to 8 months after infection (Tserel et al., 2021). It has also been found elevated in the blood plasma of subjects with chronic obstructive pulmonary disease (COPD) versus healthy controls (Acevedo et al., 2021), a disease which is associated with higher risk of hospitalization, ICU admission, and mortality from COVID-19 (Gerayeli et al., 2021). EN-RAGE has been explored in the context of COVID-19 in connection with other major inflammatory mechanisms that have been implicated in COVID-19 disease severity, including the renin-angiotensin system (Chiappalupi et al., 2021a; Chiappalupi et al., 2021b), T-cell associated cytokines (Luo et al., 2021), and a dysregulated macrophage population (Chen et al., 2022). The variety of mechanisms that EN-RAGE intersects with suggests that further experiments are needed to define its precise role in increasing COVID-19 severity.

In addition to inflammatory pathways which have been previously reported and an ongoing focus of research to target the “cytokine storm” associated with progression to ARDS (Hu et al., 2021; Song et al., 2020b; Wang et al., 2020b), our analysis also identified a cluster of growth factors and associated proteins that are significantly and specifically elevated in severe COVID-19, including FGF-5, CSF, EPHA2, TGF-α, and β-NGF. These proteins have known roles in cell proliferation of multiple tissue types, including hair growth (Higgins et al., 2014), macrophage and monocyte cell populations (Ushach and Zlotnik, 2016), lymphatic endothelial cells (Yoshimatsu et al., 2020), and the central nervous system (Iulita and Cuello, 2016), as well as various cancers (Bragado et al., 2020; Cannarile et al., 2017; Feng et al., 2020; Li et al., 2016; Li et al., 2018), suggesting a dysregulated attempt to repair tissues in response to damage caused by viral infection. Fatal COVID-19 is marked by severe pulmonary damage, diffuse alveolar damage, in conjunction with endotheliitis and microthrombosis (Bosmuller et al., 2021; Ostergaard, 2021). This damage is compounded by a dysregulated and ineffective attempt at tissue repair, with loss of basal cell populations, squamous cell metaplasia, fibrogenesis, red blood cell dysfunction and coagulation, and increased cellular senescence of epithelial and endothelial cells (D’Agnillo et al., 2021; Laforge et al., 2020). To overcome this ineffective tissue repair, several studies have suggested mesenchymal stem cell therapy as a treatment for COVID-19 (Esquivel et al., 2021; Rajarshi et al., 2020). The cell proliferation and growth markers identified in this study may have roles in regulating, and dysregulating, the process of tissue repair in COVID-19.

Of these growth markers, mesothelin (MSLN) is particularly interesting. Mesothelin is a differentiation antigen that has been almost exclusively studied in the context of malignant cancers such as mesothelioma, renal carcinoma, pancreatic cancer, ovarian cancer, and lung adenocarcinoma (Hassan et al., 2004; Pastan and Hassan, 2014), and was one of the most significantly elevated proteins in our dataset. While it is an attractive target for cancer immunotherapies due to its overexpression on cancer cells versus low expression on normal human tissue (Hassan and Ho, 2008), its function under normal physiological conditions is still largely unknown. It has been shown to bind mucin 16 (MUC16, aka CA125) (Kaneko et al., 2009), and this relationship has been suggested to contribute to metastasis (Muniyan et al., 2016); however, mouse models have shown it is not required for normal development or reproduction (Bera and Pastan, 2000), and its biological function remains ambiguous. More research is required to not only identify the mechanism by which mesothelin augments COVID-19 pathogenesis, but also the biological role of mesothelin in the lung.

Also particularly interesting are a subset of proteins significantly elevated in milder COVID-19 disease versus severe (TRANCE, FASLG, XPNPEP2, and CD207), these represent an “efficient” disease response and improved prognoses. TRANCE and FASLG were also significantly elevated in response to SARS-CoV-2 infection in subjects 18-35 years old compared to those over 65 years old. These two proteins are already known to have critical roles in regulation of the T-cell response to viral infection, and activation of the signaling pathways associated with these proteins would indicate that a robust induction of T-cell activity has taken place in response to viral infection. As mentioned above, TRANCE and FASLG are both markers T-cell activity and it is established that T-cell exhaustion is associated with more severe COVID-19 disease (Song *et al*., 2020a). TRANCE is upregulated in T-cells following antigen receptor stimulation, and promotes dendritic cell-mediated stimulation of naïve T-cells (Leibbrandt and Penninger, 2008); furthermore, mutations in TRANCE have been found to associate with higher chronicity of other viral infections (Huang et al., 2021). While upregulation has previously been reported to be associated with severe COVID-19, and chronically elevated FASLG is known to increase with aging (de Almeida Chuffa et al., 2022; Leonardi et al., 2022), none of these studies have directly compared circulating protein levels in severe cases to mild and moderate disease and chronic elevation may correlate to an inadequate T-cell response to viral infection in older people. Additionally, while these studies suggest that elevated expression of FASLG is indicative of its role in T-cell- and NK-cell-mediated apoptosis, FASLG also functions as a modulator of T-cell differentiation through non-apoptotic signaling, and facilitates the clearance of activated T-cells and B-cells cells (Nagata, 1999) as well as promoting the resolution of type 2 lung inflammation (Sharma et al., 2012; Williams et al., 2018). Therefore, a multifaceted role for elevated FASLG expression in COVID-19 is not necessarily contradictory, and this pathway has been previously proposed as a mechanism behind the abnormal T-cell activity and subsequent exhaustion observed in severe cases of the disease (Leonardi *et al*., 2022; Leonardi and Proenca, 2020). Additionally, other published studies have either only focused on mRNA concentrations and not protein (Ramljak et al., 2021), or compared “severe” ICU patients to “moderate” non-ICU hospitalized patients (Andre et al., 2022), and not to “mild” cases. The lower levels observed in severe and moderate cohorts in this study may reflect the conclusion of the process of T-cell exhaustion, while the mild cohort maintains T-cell activation and proliferation at the time of sample collection. Our data suggests that a significantly higher activation of FASLG in response to infection occurs in milder disease. Therefore, the significantly augmented levels of FASLG protein associated with severity and age suggest that circulating levels of these proteins may correlate to a more effective response to SARS-CoV-2 infection. Our data underlines the potential importance of these proteins as targets for stimulating an effective response to SARS-CoV-2 infection and require further investigation in this context.

XPNPEP2, a bradykinin-degrading hydrolase, is heavily associated with ACE activity (Rasmussen et al., 2020), and variants in the coding gene are associated with higher risk for ACE-inhibitor induced angioedema (Montinaro and Cicardi, 2020; Pall et al., 2021). Given that a “bradykinin storm” has been implicated in elevating disease severity during SARS-CoV-2 infection (Garvin et al., 2020; Roche and Roche, 2020; van de Veerdonk et al., 2020), and given XPNPEP2’s function in degrading bradykinin, elevated XPNPEP2 expression may have a potential protective effect in decreasing the inflammatory effects of bradykinin signaling, an effect supported by the data presented in this study. Due to its association with ACE, XPNPEP2 has been previously identified as potentially involved in SARS-CoV-2 infection in exploratory analyses using protein-protein interaction network databases (Khodadoost et al., 2020; Saih et al., 2021), and has been significantly associated with asymptomatic COVID-19 versus both symptomatic COVID-19 and healthy controls in pregnant women (Foo et al., 2021). However, XPNPEP2 has not yet been thoroughly investigated as a modulator of COVID-19 disease severity.

CD207, also known as langerin, is involved in efficient antigen presentation in DCs (Idoyaga et al., 2008) and has been shown to bind the SARS-CoV-2 spike protein (Gu et al., 2022), suggesting that it may play a role in viral entry, though this role has not been investigated further. However, studies showing that impairment of DC numbers and function is associated with severe COVID-19, and that this impairment can persist long after resolution of infection (Perez-Gomez et al., 2021; Zhou et al., 2020), along with observation of an increase in mature DCs in the bronchoalveolar lavage fluid of COVID-19 patients versus healthy controls (Xiong et al., 2020), support an important role for DCs in mediating COVID-19 disease response. In addition, our data supports further investigation into CD207’s interactions with SARS-CoV-2 and its role in efficient DC-mediated disease response.

In conclusion, analysis of protein concentration in plasma samples of patients infected with SARS-CoV-2 and presenting with differential severity of disease has identified proteins as biomarkers of both severe and effective responses to viral infection in mild cases. Our findings simultaneously assess a broad range of physiological responses to the disease, finding significant markers associated with multiple inflammatory pathways, viral entry and membrane fusion, endothelial damage, and cell proliferation and tissue repair, highlighting the complexity of COVID-19 pathogenesis. These proteins and associated signaling pathways are potential targets for therapeutic intervention to both stimulate an effective response to SARS-CoV-2 infection or to prevent progression to severe disease and warrant further investigation.

### Limitations of the study

Due to the patient-derived nature of the samples obtained for this study, the conclusions drawn from these results are observational, rather than mechanistic. While correlations can be made between individual protein markers that have interactions previously well-established in literature, the direct interactions between markers was not investigated in this study. Further investigation is required into the mechanistic effects of the novel protein markers highlighted in this study, for example through *in vivo* animal models of SARS-CoV-2 infection that will allow for more direct investigation.

## Acknowledgements

This research was made possible by the USC COVID-19 Biospecimen Repository, which provided patient plasma samples. This study was supported by a Keck School of Medicine (KSoM) COVID-19 research fund and the Hastings Foundation.

## Author contributions

Conceptualization, A.L.R.; Methodology, N.C.H. and A.L.R.; Investigation, N.C.H.; Formal Analysis, N.C.H.; Writing – Original Draft, N.C.H. and A.L.R.; Writing – Review & Editing, N.C.H. and A.L.R.; Funding Acquisition, A.L.R.; Supervision, A.L.R.

