## Supplemental Information for "Proteomic profiling identifies biomarkers of COVID-19 severity"

### Title

### Authors and Affiliations

Noa C. Harriott[1,2,3], Amy L. Ryan[1,2,3*]

[1]Hastings Center for Pulmonary Research, Division of Pulmonary, Critical Care and Sleep Medicine, Department of Medicine, University of Southern California, Los Angeles CA 90033

[2]Department of Stem Cell Biology and Regenerative Medicine, University of Southern California, Los Angeles CA 90033

[3]Department of Anatomy and Cell Biology, Carver College of Medicine, University of Iowa, Iowa City IA 52240

*Corresponding Author

### Contact Information

Amy L. Ryan, PhD

Previously known as Amy L. Firth

Associate Professor: Anatomy and Cell Biology
Associate Director: Center for Gene Therapy

BSB, 1-400 Core
University of Iowa
51 Newton Road
Iowa City, Iowa, 52242


### Conflict of Interest Statement

The authors have declared that no conflict-of-interest exists

### Supplemental Tables

**Supplemental Table S1: Age distribution of subjects providing plasma samples.** Mean ages were calculated for <18 years (N=0), 19-35 years (N=4), 36-50 years (N=17), 51-65 years (N=31), and >65 years (N=18). Data is expressed as mean ± S.E.M.

| Age group | Mean Age | SEM |
|---|---|---|
| <18 years | N/A | N/A |
| 19-35 years | 28.50 | 2.33 |
| 36-50 years | 42.82 | 1.31 |
| 51-65 years | 57.84 | 0.74 |
| >65 years | 71.83 | 1.13 |

**Supplemental Table S2: Significant differentially expressed proteins (DEPs) between Hispanic and non-Hispanic subjects providing plasma samples.** Data is collated from 48 Hispanic and 21 non-Hispanic subjects. One subject with unknown ethnicity is excluded from this analysis.

| Assay | OlinkID | UniProt | % change | p-value | adj. p-value |
|---|---|---|---|---|---|
| ADA | OID00560 | P00813 | 64.25 | 6.49E-04 | 0.0365 |
| TSLP | OID00497 | Q969D9 | 43.16 | 7.85E-04 | 0.0365 |
| FASLG | OID00694 | P48023 | -34.78 | 8.34E-04 | 0.0365 |
| CEACAM5 | OID00739 | P06731 | 126.87 | 9.67E-04 | 0.0365 |
| AREG | OID00728 | P15514 | 108.30 | 0.0010 | 0.0365 |
| S100A11 | OID00727 | P31949 | 51.70 | 0.0012 | 0.0365 |
| TRANCE | OID00521 | O14788 | -38.95 | 0.0018 | 0.0467 |
| EN-RAGE | OID00541 | P80511 | 95.18 | 0.0020 | 0.0467 |
| MCP-3 | OID00474 | P80098 | 145.71 | 0.0035 | 0.0637 |
| MIA | OID00701 | Q16674 | -19.46 | 0.0037 | 0.0637 |
| ANXA1 | OID00745 | P04083 | 64.72 | 0.0039 | 0.0637 |
| SYND1 | OID00664 | P18827 | 85.11 | 0.0042 | 0.0637 |
| MCP-1 | OID00484 | P13500 | 40.63 | 0.0094 | 0.1335 |
| PD-L1 | OID00518 | Q9NZQ7 | 54.68 | 0.0169 | 0.2037 |
| IL8 | OID00471 | P10145 | 57.62 | 0.0177 | 0.2037 |
| IL-18R1 | OID00517 | Q13478 | 49.00 | 0.0177 | 0.2037 |
| MSLN | OID00660 | Q13421 | 66.60 | 0.0211 | 0.2185 |
| CXCL1 | OID00496 | P09341 | 42.55 | 0.0214 | 0.2185 |
| TNFB | OID00561 | P01374 | -24.48 | 0.0282 | 0.2733 |
| CD8A | OID05124 | P01732 | -27.28 | 0.0323 | 0.2969 |
| FURIN | OID00688 | P09958 | 19.98 | 0.0353 | 0.3092 |
| IL6 | OID00666 | P05231 | 115.36 | 0.0480 | 0.3749 |

### Supplemental Figures

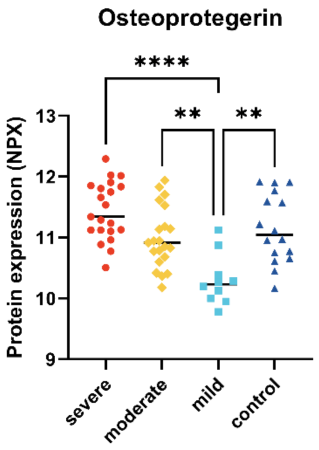


**Supplemental Figure S1: Related to Figure 4: Osteoprotegerin (OPG) expression shown by COVID-19 severity cohort.** Cluster plot of normalized protein expression (NPX) values by cohort for severe (N=21), moderate (N=21), mild (N=10), and control (N=16) cohorts. All data points are shown. **p<0.01, ****p<0.0001.

**Supplemental Database: Normalized protein expression (NPX) data provided for all samples by Olink Proteomics.** DOI: 10.6084/m9.figshare.20502948
